# Reactivating vocabularies in the elderly

**DOI:** 10.1101/216283

**Authors:** M.J. Cordi, T. Schreiner, B. Rasch

**Author notes:** **Corresponding author:** Björn Rasch, University of Fribourg, Departement of Psychology, Division of Biopsychology and Methods, Rue P.A. de Faucigny 2, 1701 Fribourg, Switzerland. The work was performed at the University of Zurich, Institute of Psychology, Department of Biopsychology. The authors declare that they have no conflict of interest.

## Abstract

Quality of memory and sleep decline with age, but the mechanistic interactions underlying the memory function of sleep in older adults are still unknown. It is widely assumed that the beneficial effect of sleep on memory relies on reactivation during Non-rapid eye movement (NREM) sleep, and targeting these reactivations by cue re-exposure reliably improves memory in younger participants. Here we tested whether the reactivation mechanism during sleep is still functional in old age by applying targeted memory reactivation (TMR) during NREM sleep in healthy adults over 60 years. In contrast to previous studies in young participants, older adults’ memories do not generally benefit from TMR during NREM sleep. On an individual level, a subgroup of older adults still profited from cueing during sleep. These improvers tended to have a better sleep efficiency than non-improvers. In addition, the oscillatory results resembled those obtained in younger participants, involving increases in theta (~6Hz) and spindle (~13 Hz) power for remembered and gained words in a later time windows. In contrast, non-improvers showed no increases in theta activity and even strongly reduced spindle power for later gained vs. lost words. Our results suggest that reactivations during sleep might lose their functionality for memory in some older adults, while this mechanism is still intact in a subgroup of participants. Further studies need to examine more closely the determinants of preserving the memory function of sleep during healthy aging.

**Grant information:** The study was supported by grant of the Swiss National Science Foundation (SNSF) No. 100014_162388. T.S. is supported by a grant of the Swiss National Science Foundation (SNSF) No. P2ZHP1_164994.

**Abbreviations:**

N1 and N2: Stage 1 and 2 sleep
SWS: Slow-wave sleep
SWA: slow-wave activity
REM: Rapid eye movement sleep
TST: Total sleep time
TMR: targeted memory reactivation

---

Aging is associated with a decrease in sleep quality. A meta-analysis of Ohayon et al. (2004) demonstrated that sleep becomes more fragmented, shorter and less deep in older adults. Sleep and particularly slow-wave sleep (SWS) is considered critical for optimal consolidation of long-term memories (e.g., Alger, Lau, & Fishbein, 2012; Diekelmann & Born, 2010; Marshall & Born, 2007). As memory formation also declines with age, several authors have proposed a link between the change in sleep quality and memory with aging (Buckley & Schatzberg, 2005; Hornung, Danker-Hopfe, & Heuser, 2005). Interestingly, according to a recent meta-analysis by Scullin and colleagues (2015), the empirical evidence for this link is rather inconsistent. While some studies reported that age-related decreases in SWS predict declines in memory consolidation (Backhaus et al., 2007; Mander et al., 2013; Westerberg et al., 2012), others observed no or even negative relationships between SWS and memory with age (Cherdieu, Reynaud, Uhlrich, Versace, & Mazza, 2014; Mawdsley, Grasby, & Talk, 2014; Mazzoni et al., 1999; Scullin, 2013). Furthermore, Wilson and colleagues reported similar benefits of sleep for memory in three age groups, in spite of strong differences in sleep quality and SWS (Wilson, Baran, Pace-Schott, Ivry, & Spencer, 2012). In contrast, several other studies report no evidence of a beneficial role of sleep compared to younger subjects (Aly & Moscovitch, 2010; Scullin, 2013). Others have suggested to include more detailed topographical information to reveal specific associations between slow-wave sleep and memory in the elderly (Mander, Winer, & Walker, 2017). Thus, the association between sleep and memory in old age is still largely unknown.

On a mechanistic level, it is widely assumed that the beneficial role of sleep for memory relies on a spontaneous reactivation of memory traces during SWS (Diekelmann & Born, 2010; Pavlides & Winson, 1989). According to the active system consolidation hypothesis, recently acquired memories are repeatedly reactivated during subsequent sleep. This reactivation is orchestrated by a fine-tuned interaction between cortical slow waves (<1 Hz), thalamo-cortical spindle activity (~13 Hz) and hippocampal sharp-wave ripple activity (100-300 Hz), and support the gradual redistribution of memories from temporary storage sites to a long-term integration in cortical memory networks. Interestingly, Gerrard and colleagues (2008) have already shown that reactivation of learned sequences during sleep is impaired in older as compared to younger rodents (Gerrard et al., 2008). Thus, one could hypothesize that the efficacy and functionality of the reactivation mechanism for memory gradually declines with age. This process could in theory be independent of the decline in general sleep quality. Alternatively, one could assume that the reactivation mechanism remains completely unaffected by age, and the impaired efficacy is solely due to the reduction in sleep quality and amount of slow-wave activity.

In this study, we started to test the functionality of memory reactivation during sleep for memory formation in older adults. In younger participants, several studies from our lab and others have now reliably established that re-exposure to memory cues during NREM sleep improves later retrieval performance (Antony, Gobel, O’Hare, Reber, & Paller, 2012; Rasch, Büchel, Gais, & Born, 2007; Schreiner & Rasch, 2015). In addition, successful memory reactivation (i.e. the difference between later remembered vs. later forgotten stimuli after targeted memory reactivation (TMR) during sleep) is characterized by specific increases in oscillatory power in the theta band (ca. 6 Hz) and spindle band (ca. 13 Hz) (Groch, Schreiner, Rasch, Huber, & Wilhelm, 2016; Lehmann, Schreiner, Seifritz, & Rasch, 2016; Oyarzún, Morís, Luque, de Diego-Balaguer, & Fuentemilla, 2017; Schreiner, Lehmann, & Rasch, 2015). According to our working model, increase in the theta band might reflect successful reinstatement of memory traces by cueing during sleep, whereas increased activity in the spindle band relates to processes of integration and stabilization of memories after reinstatement of the memory trace by the cue (Schreiner & Rasch, 2016). In the current study, we applied our established TMR paradigm using Dutch-German vocabulary to healthy participants over 60 years and examined the oscillatory correlates of successful reactivation during sleep. Based on the initial findings in rodents, we hypothesized that the benefit of reactivating memories in older adults is reduced as compared to younger participants. In addition, we expected intact theta responses during successful reactivation, indicating successful reinstatement of memories after cue re-exposure during NREM sleep. In contrast, we predicted altered spindle responses suggesting that despite successful reinstatement by the cue, memory traces are less efficiently stabilized and integrated after their reactivation in older participants.

## Materials and Method

### Subjects

A total of 23 healthy, German-speaking older adults (*n* = 8 males, ranged 62 to 83 years, mean age of 71.00 ± standard deviation [*SD*] of 5.86) took part in the experiment. None of them had intercontinental flights or shiftwork within eight weeks before participation. On the experimental day, they refrained from drinking alcohol or caffeine and got up before 8 a.m.. None had knowledge of Dutch. For participating in both sessions, participants received 120 CHF. The Ethics Committee of the Faculty of Philosophy of the University of Zurich approved the study. All participants signed a written consent prior to participation.

### Procedure

The participants were invited to two sessions which took place in the sleep laboratory. The first session started at 9:30 p.m. with some questionnaires and the attachment of the electrodes. Subjects were then allowed to go to bed and sleep until 6 a.m. This adaptation session aimed at familiarizing the subjects with the laboratory and the EEG electrodes. The second session was the experimental night and started at 9 p.m. with the same questionnaires and the EEG attachment. Before participants could go to bed, they learned 60 vocabulary pairs which were consecutively presented. Thereafter, two feedback and one final recall trial without feedback were presented, the latter measuring presleep memory performance. Half of the correctly recalled and half of the incorrect or not remembered words were randomly selected for nocturnal presentation. The volume for word replay by night was individually calibrated according to the hearing threshold test. Subjects were then allowed to fall asleep. As soon as stable NREM sleep (sleep stages N2 or SWS) appeared, reactivation was started, beginning with 5 repetitions of a test-word increasing in volume to adapt to the later reactivation volume in order to reduce the risk of awakening subjects. After a minimum of 3 hours of sleep and 90 minutes of reactivation, subjects were awakened to recall all the words again. Again, no feedback was provided. Thereafter, they were allowed to return to bed and sleep until 6 a.m. (see Figure 1A for the procedure of the experiment).

**Figure 1.**
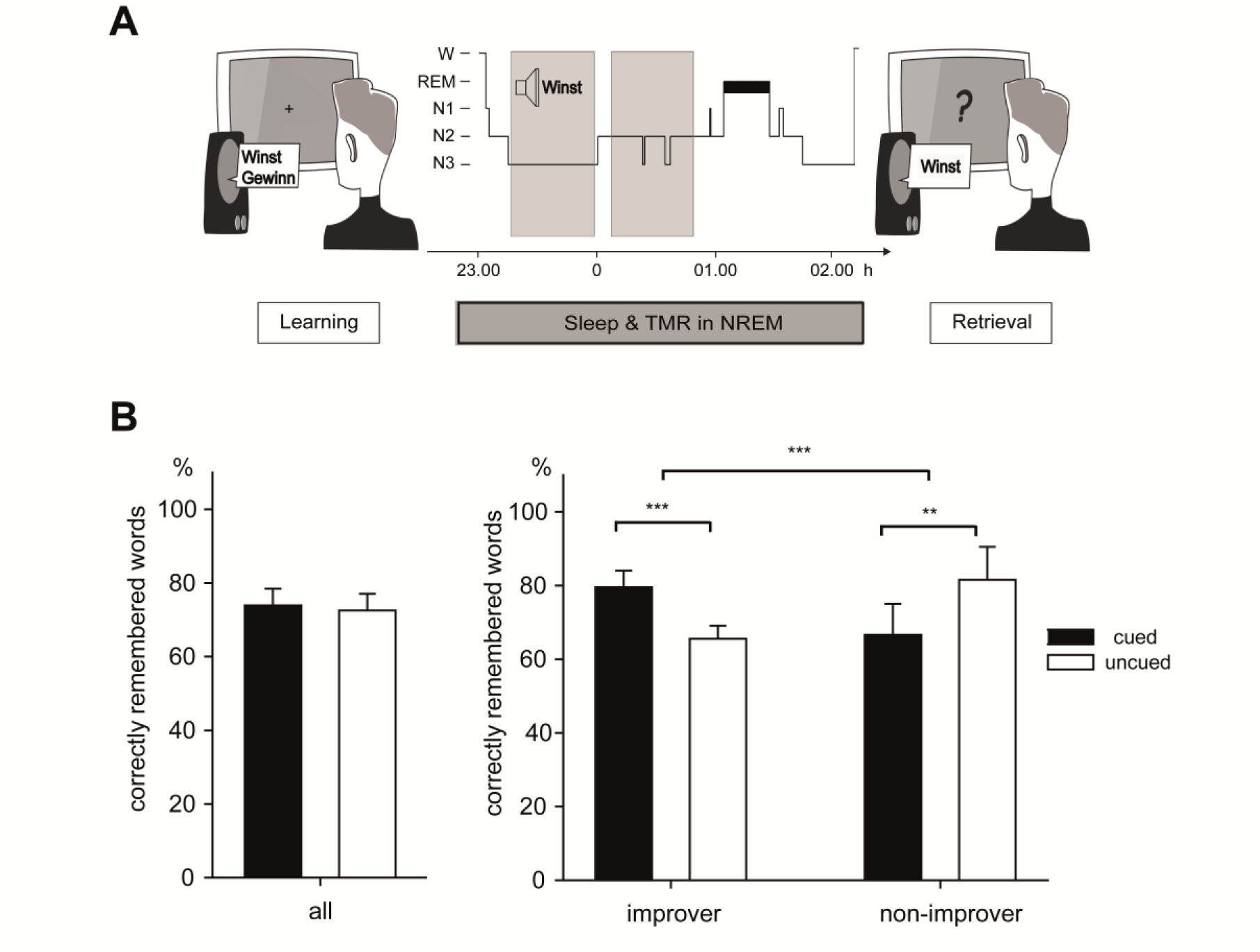
Experimental procedure and behavioral results. (A) The experimental procedure (reproduced from Schreiner & Rasch, 2015) and (B) the behavioral results over all as well as separately for improvers and non-improvers are displayed. Subjects learned 60 Dutch-German word pairs before sleeping. During the retention interval, 50% of the correctly recalled and 50% of the not or incorrectly remembered words were replayed during NREM sleep (TMR). After 90 minutes of reactivation, participants were tested on their memory for the word pairs in a cued recall design. Results showed that subjects did not generally benefit from TMR (left panel). Due to huge individual differences, we split the sample into “improvers” who benefitted from cueing and “non-improvers” who did not (right panel). Asterisks indicate level of significance ** *p* < .01, *** *p* < .001.

### Hearing Threshold Level Test

To individually adapt the nightly volume of replayed vocabulary, a hearing threshold level test was done. A test-word was repeatedly presented over the loudspeakers which were placed in a similar distance to the subject as during later sleep. The volume of the word was regulated down and participants were asked to increase the volume until they could hear the voice. The loudness was measured with a decibel measure and the value was noted. Then, the participants were asked to reduce the volume until he could hardly hear the spoken word. Again, volume was measured and noted. This procedure was repeated three times. The mean from these six values was taken as hearing threshold level. During the experimental night, the words were presented with a volume of 15 dB above this threshold. Mean volume with which the words were presented was 71.16 ± 5.58 dB.

### Vocabulary Task

We used the vocabulary learning task as described by Schreiner and Rasch (2015). We adapted the difficulty of the task by reducing the word list to 60 and by including a second feedback learning trial. The Dutch words were presented acoustically via loudspeakers, followed by a fixation cross on the screen (500 ms). Subsequently, the corresponding German translation was presented in black font on the white screen for 2000 ms. A blank screen separated the trials (2000 – 2200 ms). Subjects were asked to memorize as many words as possible. In a second and third learning trial, a question mark appeared instead of the German translation after the spoken Dutch word and the fixation cross. Subjects were asked to name the corresponding translation or to say “next” in case they did not remember the word. Feedback was given by presenting the correct answer. The fourth iteration, in which no feedback was provided, was taken as presleep recall performance. The learning performance level before sleep was on average 61.02% out of the 60 words, indicating a medium task difficulty, excluding ceiling or floor effects.

After a minimum of 90 minutes of reactivation, subjects were awakened and recall was again tested by presenting the Dutch word and asking the participant to give the German translation. Again, no feedback was provided. The order in which the words were recalled was different across all three recall phases. The relative performance level of postsleep and morning recall was measured by setting presleep performance to 100%.

### Reactivation of Vocabulary

After the last learning trial, an automatic MATLAB algorithm randomly extracted 50% of all correctly recalled and 50% of all incorrect or not recalled words. Thus, 30 words were individually chosen for replay during later sleep. Only the Dutch words, without their German translation were presented via loudspeakers during sleep. During reactivation, one word appeared every 3.8 – 4.2 ms in a randomized order, resulting in on average 33.7 ± 6.34 exposures per word. Sleep was scored online and reactivation was immediately stopped as soon as indications of REM sleep, arousals or awakenings appeared. Postexperimental offline sleep scoring confirmed that on average 98.22 ± 1.67% of the words were correctly presented during N2 or N3.

### Polysomnographic Recordings

Electromyographic (EMG), electrocardiographic (ECG), electrooculographic (EOG), and electroencephalographic (EEG) electrodes were attached for polysomnography. Impedances did not exceed 80 kΩ. High density EEG was recorded with a 128 channels Geodesic Sensor Net (Electrical Geodesics, Eugene, OR) and a sampling rate of 500 Hz. For sleep scoring, data was filtered according to the settings suggested by the American Association of Sleep Manual (AASM). Those include a low frequency filter at 0.3 Hz and a high frequency filter at 35 Hz for the EEG electrodes. The EMG signal was filtered between 10 and 100 Hz. Two independent sleep scorers visually scored 30 second segments of sleep to define the stages 1-3, REM sleep, and wakefulness offline and according to standard criteria (Iber et al. 2007). Derivations F4, C4, O4, HEOG, VEOG, and EMG were used therefore. A third sleep expert was consulted in case of disagreement. For further analyses, segments with sleep stages 2 and 3 were included only.

### EEG Data Analysis

Preprocessing was done with Brain Vision Analyzer 2.1 (Brain Products, Gilching, Germany). The electrodes were re-referenced against the mean of the mastoids (electrodes 100 and 57) and filtered (low-pass filter: 0.1 Hz; high pass filter: 35 Hz). Only segments scored as N2 and N3 were selected in order to exclude possible reactivations administered in a wrong sleep stage. Segments were built within -1000 to 4000 ms around the cues. Artefacts were excluded semi-manually. Voltage differences larger than 400 µV within an interval of 200 ms were automatically detected. Those were afterwards screened and deleted manually.

#### Event-Related Potentials (ERP) analysis

An interval of -500 ms to -100 ms before the stimulus onset was taken as baseline and used for baseline correction. For each subject, ERPs were averaged for each condition: overnight gains (words not remembered before, but after sleep), losses (words remembered before, but not after sleep), hits (words remembered before and after sleep) or misses (words remembered neither before not after sleep), remembered (hits + gains) and forgotten (misses + loss) words. As in Schreiner and Rasch (2015), the average EEG amplitudes in the interval from 800 to 1100 ms after the stimulus were exported to SPSS for statistical analysis.

#### Power-analyses

Power-analyses were performed with Brain Vision Analyzer on 2 second segments of NREM sleep (1024 data points, 10% overlap) using a Fast Fourier Transformation (FFT) with a Hamming window of 10%. We exported the area information of slow-wave activity (SWA, 0.5-4.5 Hz), theta activity (4.5-8 Hz), the slow spindle band (11-13 Hz) and fast spindle band (13-15 Hz) into SPSS to run group-wise comparisons on frontal, central and parietal left and right electrodes to compare overall power band differences.

#### Wavelet-analysis

The wavelet analyses were performed with fieldtrip (Oostenveld, Fries, Maris, & Schoffelen, 2011) after preprocessing. Here, condition based segments were created according to overnight gains, losses, hits or misses, remembered (hits + gains) and forgotten (misses + losses) words, each beginning 1000 ms before the cue and ending 4000 ms after the reactivation. As not each participant had gained or lost words, the sample in which we could analyze gains and losses was reduced (*n* = 15) compared to the main sample (*n* = 23). We selected a time window of -500 to -100 ms before the reactivation cue that served as the baseline for the calculation of the relative power change. The values thus represent power changes ranging from 0 to 1. For statistical analyses, these relative power values were exported to SPSS for the investigated frequency and time ranges. In a first step, data was analyzed according to the time and frequency windows reported in Schreiner et al. (2015). Secondly, post hoc explorative analyses were conducted according to visual inspection of the time-frequency plots (see Figures 2 D, 2 F, Figure 3 and Figure 4 E – H).

**Figure 2.**
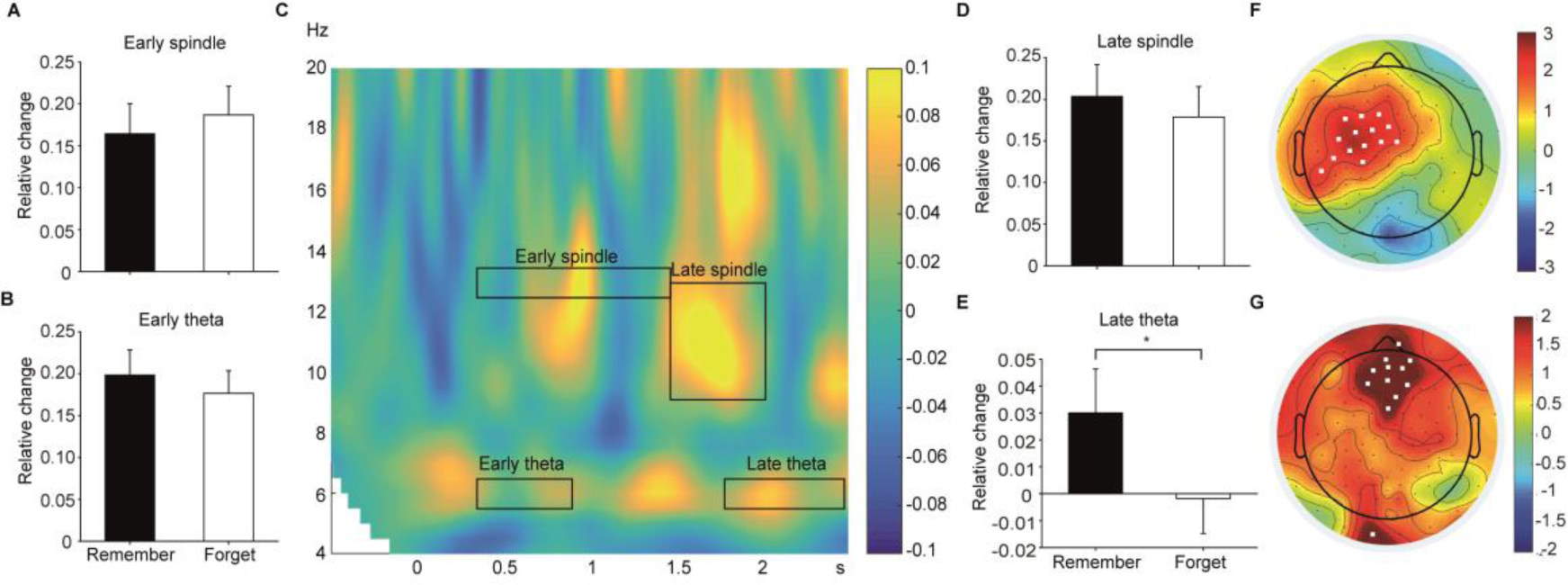
Main results. The bar plots display the main effect of cueing in (A) Early spindles (12.6-13.6 Hz, 400 – 1500 ms), (B) Early theta (5.6-6.6 Hz, 400 – 900 ms), (D) Late spindle (9.6-13 Hz, 1500 – 2100 ms) and (E) Late theta (5.6-6.6 Hz, 1800 – 2500 ms). Asterisks indicate significant results * *p* < .05. A relative difference of 0.1 points corresponds to 10% change. (C) The time-frequency analysis across all subjects was performed exemplarily in electrode Fz. The four investigated areas are indicated by a black square. The areas of early theta/spindle and late theta were a priori defined based on Schreiner et al., 2015. The area for late spindle represents an exploratory analysis. The topoplots display t values of remembered versus forgotten words in all subjects in Late spindle (F) and Late theta (G). Significant electrodes are marked in white colour (*p* < .05, uncorrected).

**Figure 3.**
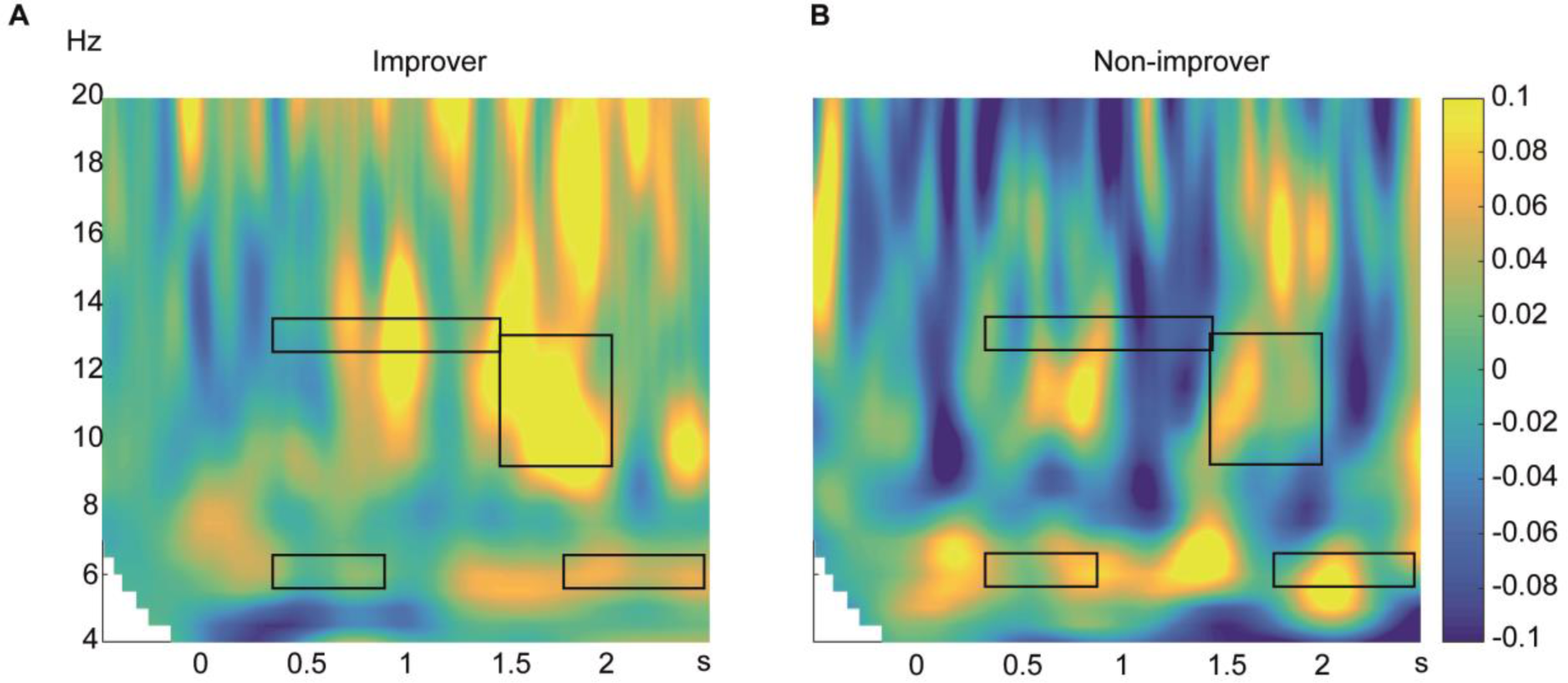
Time-frequency analysis separately for improvers and non-improvers. Time-frequency analyses in improvers (A) and non-improvers (B) separately for the four investigated areas on remembered versus forgotten words. As example electrode we display Fz here.

**Figure 4.**
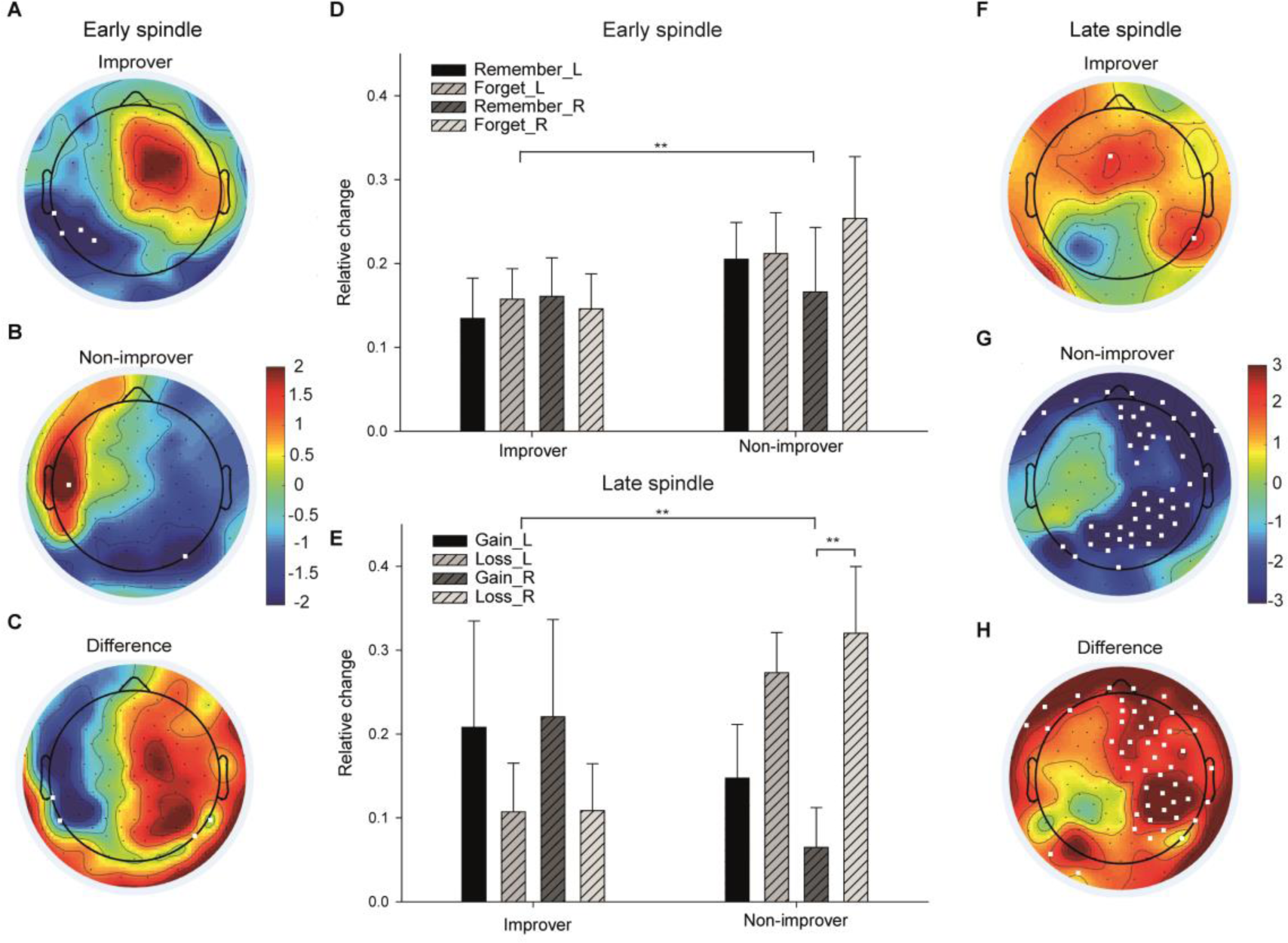
Results for Early and Late spindle. Topoplots A – C show t-values of remembered versus forgotten words in improvers (A), non-improvers (B), and the difference between those two groups (C) in the Early spindle band (12.6-13.6 Hz, 400 – 1500 ms). Significant electrodes are marked in white colour (*p* < .05, uncorrected). Plot D displays the according bars separately for left and right hemisphere. Topoplots F – H show the t-values of gained versus lost words in the late spindle band (9.6 – 13 Hz, 1500 – 2100 ms). Its according barplot is segment E. Asterisks indicate significances ** *p* < .01.

### Statistical Analysis

The data was analyzed using analyses of variance with the between subjects factor group (improver vs. non-improver) and the within subjects factor cue (remember vs. forget or gain vs. loss, respectively). In additional analyses we also included the within-subjects factors hemisphere (left vs right) and topography (frontal vs central vs parietal). For follow-up analyses, paired-samples t-tests within the groups or t-tests for independent samples were used. Values were adapted when equal variances could not be assumed. The level of significance was set to *p* = .05 (uncorrected).

## Results

### Behavioral Data

In contrast to our previous findings in younger adults (Schreiner et al., 2015; Schreiner & Rasch, 2015), presentation of single Dutch words during sleep did not improve retrieval performance of the German translation in older adults: Participants remembered 73.93 ± 4.57% (*SEM*) words that were replayed during sleep (“cued” words) and 72.52 ± 4.56% uncued words, with memory performance before sleep set to 100%. Memory for cued and uncued words did not significantly differ (*p* > .70) (see Figure 1B). Generally, retrieval success was lower in older adults (73.22 ± 4.16%) as compared to our previous results in younger participants (94.05 ± 1.64%, Schreiner, Lehmann, & Rasch, 2015). Learning performance before sleep was generally better than in younger adults (61.02 ± 3.49% vs 51.61 ± 2.69%, respectively). Note that the task encompassed only 60 word-pairs in older adults as compared to 120 word-pairs in younger participants. Evening learning performance between cued and uncued words did not differ in the current sample (*p* ≥ .09).

In spite of the lack of an effect in older participants, the variability of cueing benefits on memory was remarkable: While some participants profited from cueing during sleep with up to 33% of memory improvement, others showed an impaired performance induced by cueing of -42%. To further explore possible systematic differences between older adults that profited from cueing and others that did not, we split our sample into one group of “improvers” (*n* = 13) and one group of “non-improvers” (*n* = 10) defined by cueing benefits above or below the overall mean (i.e. 1.41% improvement). Obviously, improvers remembered more cued than uncued words (79.56± 4.49% vs. 65.55 ± 3.52%; *t*(12) = 5.47, *p* < .001), while in non-improvers the pattern was reversed (cued vs. uncued words: 66.61 ± 8.47% vs. 81.57 ± 8.91%, *t*(9) = -3.89, *p* = .004) (see Figure 1B). Consequently, the interaction between group and cueing was highly significant (*F*(1, 21) = 42.30, *p* < .001, eta^2^ = .67). Improvers and non-improvers did neither differ in age (*p* > .80) nor in pre-sleep learning performance (*p* > .30), nor in the amount of cueing (*p* > .20).

### Sleep Data

Interestingly, older adults who profited from cueing during sleep showed a statistical trend for higher sleep efficiency (0.86 ± 0.02%) as compared to non-improvers (0.79 ± 0.03%; *p* = .10). In particular, improvers fell asleep faster than non-improvers (7.92 ± 2.64 min vs. 18.35 ± 5.36 min, *p* = .10). Descriptively, they also spent more time in NREM sleep and slow-wave sleep, although these differences did not reach significance (228.04 ± 13.54 min vs. 194.95 ± 17.73 min NREM, *p* > .15 and 74.19 ± 14.47 min vs. 50.55 ± 10.66 min SWS, *p* > .23, respectively). Similarly, no other sleep stage did significantly differ between the two groups (all *p* > .30, see Table 1). We also compared overall power differences in several other frequency bands between improvers and non-improvers in NREM sleep. We found no main effects of group for fast spindles (13-15 Hz), *p* > .90, slow spindles (11-13 Hz), *p* > .90, theta activity (4.5-8 Hz), *p* > .70 or slow-wave activity (0.5-4.5 Hz), *p* > .28, during sleep. Also, no significant correlation of the cueing benefit on memory with any sleep stage or general power was observed, neither on the overall group level nor in the separate groups of improver and non-improvers (all *p* > .10).

**Table 1.** Sleep stages during cueing in improvers and non-improvers

| Sleep stage | Improvers (n=13) | Non-improvers (n=10) | *p* |
| --- | --- | --- | --- |
|  | *M* ± *SEM* | *M* ± *SEM* |  |
| % Wake | 12.07 ± 2.02 | 15.28 ± 2.44 | .32 |
| % N1 | 5.51 ± 0.80 | 6.96 ± 1.26 | .32 |
| % N2 | 50.60 ± 5.10 | 53.24 ± 4.27 | .71 |
| % N3 | 26.56 ± 6.03 | 18.64 ± 3.60 | .31 |
| % NREM | 77.16 ± 2.41 | 71.87 ± 2.92 | .17 |
| % REM | 5.21 ± 1.22 | 5.82 ± 2.06 | .79 |
| Sleep latency (min) | 7.92 ± 2.64 | 18.35 ± 5.36 | .10 |
| SWS latency (min) | 22.69 ± 5.82 | 34.90 ± 10.91 | .31 |
| REM latency (min) | 170.42 ± 26.76 | 164.30 ± 24.03 | .87 |
| Total min | 299.23 ± 20.38 | 272.30 ± 22.68 | .39 |
| Sleep efficiency | 0.86 ± 0.02 | 0.79 ± 0.03 | .10 |

*Note.* Sleep stages in % or minutes ± standard errors of the mean (SEM) separately for improvers and non-improvers. P-values indicate group comparisons. Sleep efficiency was calculated as minutes in N1, N2, N3 and REM divided by the sum of total minutes asleep and sleep latency.

### ERP Analysis

The average ERP amplitudes of later remembered versus forgotten words did not differ (all *p* ≥ 1 after Bonferroni-correction).Neither was there any difference between lost and gained words (all *p* > .04).

### Wavelet Analysis

To analyze the oscillatory response to cues during sleep in older adults we explored the difference between successful and unsuccessful cueing in the time-frequency space. In a first step, we restricted our analysis to frequency bands and time intervals previously reported for younger adults (see Figure 2): For theta activity (~ 6 Hz), we used an early time window (400 – 900 ms) and a late time window (1800 – 2500 ms). For spindle activity (~ 13 Hz) we used one time window between 400 and 1500 ms (see Schreiner, Lehmann & Rasch, 2015, for details). As effects in younger adults had been particularly reported in frontal regions, electrodes were grouped in six topographical regions (frontal/central/parietal x left/right) and included as within-subject factors in an ANOVA as done previously (see Cordi, Schlarb, & Rasch, 2014 for details).

In contrast to results in younger participants, older adults did not show any significant increase in theta activity in the early time interval after the reactivation cue (400 – 900 ms), neither on the general level nor for any group differences (all *p* > .20) (see Figure 2B). However in the late time window (1800 – 2500 ms) of younger participants, older adults also showed generally increased theta activity for later remembered as compared to forgotten words during sleep (*F*(1, 21) = 4.34, *p* = .05, eta^2^ = .17), (see Figure 2E and G). No interactions with the topography or lateralization were observed. Descriptively, improvers showed more robust theta activity in this late time window over mid- and right-frontal regions (see Figure 2 C-E). However, we did not find any significant differences between the two groups (all *p* > .60). However, when focusing the analysis on “gained” words by cueing (cued words not remembered during learning, but after sleep) and “lost” words (cued words remembered during learning, but not after sleep), a significant three-way interaction of topography, cue and group occurred (*F*(2, 26) = 5.561, *p* = .010, eta^2^ = .30). Particularly in frontal electrodes, improvers had significantly higher theta power after gains (.226 ± .078) than non-improvers (-.068 ± .045, *t*(13) = -2.861, *p* = .013), while average theta activity did not differ significantly for losses between the two groups (*p* = .475). In addition, the main effect of successful cueing in the late theta window was confirmed also in gained vs. lost words (*F*(1, 13) = 5.48, *p* = .036, eta^2^ = .297). Please note that in comparison to younger adults, the level of gains was generally much lower in older adults. One of the 23 subjects had no losses while 7 had no gains at all and had to be excluded from this analysis. Therefore, the analysis on gains and losses encompasses only 15 subjects (9 improvers, 6 non-improvers).

With respect to increases in spindle power previously observed in younger participants (~ 13 Hz, time window between 400 and 1500 ms), older subjects rather showed a reduction in spindle power to later remembered as compared to later forgotten cued words (see Figure 2A): particularly on parietal recording site, spindle power was generally lower to remembered (0.123 ± 0.04) as compared to forgotten words (0.174 ± 0.03; *t*(22) = -2.032, *p* = .054). No effect occurred on other electrode sites (all *p* ≥ .70, topography x cue interaction: (*F*(2, 42) = 4.078, *p* = .024, eta^2^ = .16). Interestingly, tendencies for spindle power increases and decreases are exactly on opposite hemispheres in improvers and non-improvers. Improvers and non-improvers thus differ in their spindle power pattern in both hemispheres (see Figure 4 A – C). This is substantiated by a significant three-way interaction between hemisphere, cue, and group (*F*(1, 21) = 6.74, *p* = .017, eta^2^ = .24) (see Figure 4 D). With respect to gains vs. losses in the spindle band the topography*cue interaction as well as the hemisphere* cue* group interaction observed for remembered versus forgotten words appeared as a trend (*p* = .076 and *p* = .066, respectively). No other comparisons were significant (all *p* > .20).

However, post-hoc visual inspection of the data revealed that the spindle result pattern was even more pronounced in a later time-window between 1500 - 2100 ms and extended also to slower spindle frequencies (9.6 - 13 Hz) (see Figure 2D for remembered vs. forget and Figure 4 E - H for gains vs. losses). While improvers descriptively had still increased spindle power (gains: 0.21 ±.12 vs. losses: 0.11 ± 0.06), *t*(8)= 1.36, *p* = .21), non-improvers exhibited a strong reduction in spindle power for gains (0.11 ± 0.05) as compared to losses (0.30 ± 0.06), *t*(5) = -5.47, *p* = .003). The interaction between cue and group was highly significant (*F*(1, 13) = 8.67, *p* = .011, eta^2^= .40) while the main effect of successful cueing in the interval was not (*p* = .42).

For remember vs. forget in this frequency range (9.6-13Hz, 1500-2100ms), a main effect of topography (*F*(2, 42) = 13.45, *p* < .001, eta^2^ = .39) and the interaction between cue and topography (*F*(2, 42) = 4.37, *p* = .019, eta^2^ = .17) were significant. Follow-up t-tests revealed no significant differences between remembered vs. forgotten words in frontal, central, or parietal electrodes (*p* > .08), (see Figure 2 D and F). However, this exploratory analysis needs to be interpreted with caution, and might inspire future studies to examine slow spindle power in this later time window in cueing studies in the elderly.

## Discussion

Here we demonstrate that reactivations do not generally increase memory performance in older adults contrary to what has been reported for young adults (Schreiner & Rasch, 2015). Retrieval performance for words that were not reactivated during sleep did not differ from memory for those words that we represented during NREM sleep. Also on the oscillatory level, the characteristic increases in theta and spindle power during an early time window (0.5 – 1.5 seconds) reported in young adults were absent in older adults. Here, even decreases in spindle power associated with successful cueing were observed. Only at later time windows (> 1.5 seconds), older adults generally also had increases in theta and spindle power associated with successful cueing during NREM sleep.

Several reasons could explain the lack of positive effects of TMR during NREM sleep in older adults. First, our results could indicate that the spontaneous memory reactivation mechanism occurring during NREM sleep in younger age is less functional in old age. This explanation is supported by findings from Gerrard and colleagues (2008), who show that spontaneous sequence reactivation during NREM sleep is markedly impaired in aged as compared to young rats. In addition, spatial memory score of the animals correlated with the degree of sequence reactivation in this study. In this case, one could assume that also targeting this mechanism by external memory cues during NREM sleep is less effective in older as compared to younger individuals, possibly explaining our null findings reported here. The notion of a less functional reactivation mechanism during NREM sleep in older age is consistent with studies reporting no or only small benefits of sleep on memory in older adults: For example, Scullin and colleagues (2017) recently reported no effects of an afternoon nap on memory performance in older adults (58 – 83 years). In contrast, younger adults showed the well-known sleep-benefits on memory. Similar results have been obtained by other groups (e.g., Cherdieu et al., 2014; Scullin, Fairley, Decker, & Bliwise, 2012; Wilson et al., 2012). Conversely, several studies have shown that sleep disruption or sleep deprivation results in fewer memory impairments in older adults as compared to young one (Bonnet, 1989; Duffy, Willson, Wang, & Czeisler, 2009; Nesthus, Scarborough, & Schroeder, 1998). These data suggest that the importance of sleep for memory is markedly reduced with age, possibly due to a reduction of an effective memory reactivation mechanism during NREM sleep.

Another possible explanation for the lack of TMR-induced memory benefits is the well-known reduction in SWS with age. Some authors have reported positive correlations between memory scores and the amount of SWS (Backhaus et al., 2007; Deak, Stickgold, Pietras, Nelson, & Bubrick, 2011), possibly even mediated by the degree of prefrontal atrophy (Mander et al., 2014). However, others have reported also no or negative correlations between the amount of SWS and memory in old age (Mazzoni et al., 1999; Scullin, 2013). In the current study, older adults spent on average 64 min in deep slow-wave sleep scored by standard criteria, which appears to be sufficient for inducing memory benefits during sleep. Thus, we consider it unlikely that the amount of SWS alone can explain the lack of TMR benefit on memory in our study and the reduced importance of sleep in general for memory in older age reported by others.

We also need to consider additional differences between the current TMR study and previous studies. For example, we adapted the learning task by reducing the amount of learning stimuli (i.e., from 120 to 60 word-pairs) and by increasing the number of repetitions (i.e., from 2 to 3 learning rounds). Some studies indicate that the number of stimuli (Feld & Born, 2017) and learning ability (Tucker & Fishbein, 2008) can influence the effect of sleep on memory. In addition, Creery and colleagues (2015) showed that targeted memory reactivation does not further strengthen memories for items learned almost perfectly. However, the immediate learning performance in the current study was at 60%, which is considered an optimal immediate performance level for obtaining sleep-benefits on memory (Diekelmann, Wilhelm, & Born, 2009; Drosopoulos, Schulze, Fischer, & Born, 2007). Still, it might be possible that age-related differences in learning capabilities and encoding strategies might be the reason for the lack of sleep and TMR-induced memory benefits (Shing, Werkle-Bergner, Li, & Lindenberger, 2008).

Despite lacking a benefit on the overall level, we had observed marked inter-individual behavioral differences concerning cueing effects of 42% decrease up to 33% performance increase. Thus, we split the sample into a group of improvers and non-improvers based on whether subjects had experienced a memory boost from cues or not. It might be that improvers preserved the mechanism and rather show similarities to the pattern observed in younger adults while non-improvers suffer from age-related changes that have already happened. One could argue that this split might have resulted in an arbitrary separation, not necessarily implicating an underlying functional difference. However, for arbitrary groups we would not have expected to find any systematic differences in electrophysiological measures. Moreover, studies indicate that some older adults profit more from sleep than others do and can even relate that to biomarkers such as frontal grey matter atrophy or white matter structure in the brain (Mander et al., 2013; Mander, Zhu, et al., 2017). We unfortunately have not included such measures of brain morphology. However,, improvers appeared to have slightly better sleep efficiency than non-improvers. In addition, the oscillatory pattern was indeed systematically different in improvers and non-improvers. Importantly, the time-frequency results for improvers were more similar to those observed in young participants. For example, in the late spindle band, improvers showed significant increases in spindle power during successful cueing whereas non-improvers exhibited significant reductions. A similar pattern emerged in the early spindle band, with some increase in spindle power particularly in the right hemisphere for improvers and reductions of spindle power in non-improvers. In the theta band, the differences between improvers and non-improvers were less pronounced, with only a descriptive increase in theta power for improvers vs. non-improvers. According to our working-model (Schreiner & Rasch, 2016), increases in theta power after a memory cue during NREM sleep indicate successful reinstatement of the associated memory trace. However, successful reinstatement alone is not sufficient to achieve memory benefits by TMR. In addition, increases in spindle activity are necessary to allow a successful stabilization and integration of the reinstated memory trace. According to this model, the early reinstatement of the memory trace observed in younger participants (0.5 – 1.5 seconds) is absent in older adults, possibly reflecting the impaired functionality of the reactivation mechanism in older age. Thus, plastic processes associated with early spindle power increases might not be capable of stabilizing the memory trace, as it was not yet successfully reinstated. Also in younger adults, a recent study shows that later spindle responses between 1.7 and 2.1 seconds might be even more critical for successful consolidation processes after reactivation during sleep, as they can contain category-specific memory information (Cairney, Guttesen, El Marj, & Staresina, preprint). In the older participants, the general late theta responses indicates that successful reinstatement is achieved in a later period (> 1.5 seconds), possibly similarly in both improvers and non-improvers. Then, only in improvers, a late spindle power increase in a slower frequency band (9.6-13 Hz) occurs, which might underlie the observed memory benefit by TMR during sleep in this group. However, this speculation requires further experimental confirmation in larger samples.

In sum, our results demonstrate that external reactivation of recently learned memory content does not generally help to improve memory in older adults. These findings indicate that the mechanism underlying sleep’s support for memory consolidation changes across age. This change however seems to be subjected to inter-individual differences, so that some individuals can maintain the capabilities of memory reactivation during age even in old age. Future studies are needed to identify possible mechanisms and biomarkers for maintaining sleep’s role for memory also in old age.

## Acknowledgements

We are grateful for the support of Jasmin Widmer, Maya Thalia Schenker and Amela Kujevic in data collection. This work was funded by a grant from the Swiss National Science Foundation (SNSF) No. 100014_162388. T.S. is supported by a grant of the Swiss National Science Foundation (SNSF) No. P2ZHP1_164994.

